# The total and active bacterial community of the chlorolichen *Cetraria islandica* and its response to long-term warming in sub-Arctic tundra

**DOI:** 10.1101/2020.03.04.976944

**Authors:** Ingeborg J. Klarenberg, Christoph Keuschnig, Denis Warshan, Ingibjörg Svala Jónsdóttir, Oddur Vilhelmsson

## Abstract

Lichens are traditionally defined as a symbiosis between a fungus and a green alga and or a cyanobacterium. This idea has been challenged by the discovery of bacterial communities inhabiting the lichen thalli. These bacteria are thought to contribute to the survival of lichens under extreme and changing environmental conditions. How these changing environmental conditions affect the lichen-associated bacterial community composition remains unclear.

We describe the total (rDNA-based) and potentially metabolically active (rRNA-based) bacterial community of the lichen *Cetaria islandica* and its response to long-term warming using a 20-year warming experiment in an Icelandic sub-Arctic tundra. 16S rRNA and rDNA amplicon sequencing showed that the orders Acetobacterales (of the class Alphaproteobacteria) and Acidobacteriales (of the phylum Acidobacteria) dominated the bacterial community. Numerous ASVs (amplicon sequence variants) taxa could only be detected in the potentially active community but not in the total community. Long-term warming led to increases in relative abundance on class, order and ASV level. Warming altered the relative abundance of ASVs of the most common bacterial genera, such as *Granulicella* and *Endobacter*. The potentially metabolically active bacterial community was also more responsive to warming than the total community.

Our results suggest that the bacterial community of the lichen *C. islandica* is dominated by acidophilic taxa and harbors disproportionally active rare taxa. We also show for the first time that climate warming can lead to shifts in lichen-associated bacterial community composition.

## 1 Introduction

The notion that lichens harbour diverse bacterial and fungal communities has challenged the traditional view of the lichen as a symbiosis between a fungus (mycobiont) and an alga and/or a cyanobacterium (photobiont) (González et al. 2005; Spribille et al. 2016). Nonetheless, the first lichen-associated bacteria were already discovered in the 1920s (Uphof 1925). To date, bacterial communities of a wide-range of lichen species have been revealed by molecular approaches (Cardinale, Puglia, and Grube 2006; Cardinale et al. 2008; Grube et al. 2009; Hodkinson and Lutzoni 2009; Bjelland et al. 2011; Bates et al. 2011; Mushegian et al. 2011; Sigurbjörnsdóttir, Andrésson, and Vilhelmsson 2015; Park et al. 2016). Alphaproteobacteria usually dominate the lichen microbiome, but other taxa such as Actinobacteria, Firmicutes, Acidobacteria, Betaproteobacteria, Deltaproteobacteria and Gammaproteobacteria are also found. These bacteria can form highly structured, biofilm-like assemblages on fungal surfaces and within the lichen thallus (Grube et al. 2009). The bacterial communities inhabiting the lichen thalli play important roles in the lichen holobiont (the lichen and its microbiome), by contributing to nutrient supply, resistance against biotic and abiotic stresses, production of vitamins and support of fungal and algal growth by the production of hormones, detoxification of metabolites and degradation of senescing parts of the lichen thallus (Sigurbjörnsdóttir, Andrésson, and Vilhelmsson 2015; Grube et al. 2015). Thereby, the bacterial part of the lichen holobiont is suggested to contribute to the survival of lichens under extreme and changing environmental conditions.

The composition of associated bacterial communities of lichens may be shaped by intrinsic and extrinsic factors. Among intrinsic factors affecting the lichen microbiome are thallus age (Cardinale, Steinová, et al. 2012), mycobiont species and photobiont species (Coleine et al. 2019; Hodkinson et al. 2012; Grube et al. 2009; Bates et al. 2011; Wedin et al. 2016). The composition of lichen bacterial communities can also be influenced by extrinsic factors such as sunlight exposure and substrate type (Cardinale, Steinová, et al. 2012; Park et al. 2016), geography and local habitat (Cardinale, Grube, et al. 2012; Printzen et al. 2012; West et al. 2018; Hodkinson et al. 2012), altitude (Coleine et al. 2019), drought (Cernava et al. 2019) and arsenic contamination (Cernava et al. 2018). While some lichens can adapt to changing environmental factors by switching photobionts depending on the ecological niche of the photobiont (Rolshausen et al. 2018; Domaschke et al. 2013), not all lichens are able to adapt to changing environments in this way. Lichens might also be able to acclimate through changes in their associated bacterial communities. This strategy has been demonstrated for several environmental factors, such as drought (Cernava et al. 2019) and arsenic contamination (Cernava et al. 2018). Substrate type is another extrinsic factor that can influence the composition of bacterial communities. Therefore, changes in C (carbon) or N (nitrogen) availability in the environment, for instance as a result of changes in plant litter quality due to shrubification, might be factors altering the structure of the lichen microbiome. However, little is known about the effect of long-term environmental changes on the bacterial communities associated with lichens.

High-latitudes are especially rich in lichen species and biomass (Nash 2008; Cornelissen et al. 2007), where they make significant contributions to ecosystem functioning (Asplund and Wardle 2017). Mat-forming lichens such as Cetraroid species contribute to primary production and nutrient cycling, control soil chemistry and water retention (Cornelissen et al. 2007). Currently, climate in high-latitudes warms twice as fast as elsewhere (IPCC 2019) resulting in increased abundance of shrubs, particularly in the low and sub-Arctic (Elmendorf et al. 2012; Myers-Smith et al. 2019). Direct effects of warming on lichens include changes in C-based secondary compounds (Asplund, Siegenthaler, and Gauslaa 2017) and increased biomass (Biasi et al. 2008). Warming also has indirect effects on lichens. In many low and sub-Arctic tundra ecosystems shrubification results in increased shading and greater amounts of litter, which can lead to decreased lichen photosynthesis rates causing a decline in lichen biomass (Elmendorf et al. 2012; Cornelissen et al. 2001; Fraser et al. 2014; Alatalo et al. 2017; Nash and Olafsen 1995). Yet, the effect of long-term warming on the bacterial communities of lichens in high-latitudes needs to be investigated.

In this study we investigate the total and potentially metabolically active bacterial community of the lichen *Cetraria islandica* (L.) Ach. (English ‘Iceland moss’) and its response to two decades of warming in open top chambers (OTCs) in an Icelandic sub-Arctic alpine dwarf-shrub heath. *C. islandica* is a mat-forming chlorolichen with foliose thalli and forms a major component of the vegetation in Arctic, sub-Arctic and alpine environments throughout the northern hemisphere (Kärnefelt, Mattsson, and Thell 1993). 16S rRNA and rDNA sequencing was used to characterize the potentially active and total bacterial community in control plots and OTCs. We also quantified 16S rRNA gene abundance by quantitative PCR to compare the absolute abundance of bacteria in the controls and OTCs. Finally, it was recently demonstrated that N_2_-fixing bacteria could associate with chlorolichens (Almendras et al. 2018). Thus, we also quantified the number of *nifH* genes by quantitative PCR in order to test if the *C. islandica* microbiome could potentially perform N_2_-fixation and how warming influences the abundance of associated N_2_-fixers.

We predicted that long-term warming and the associated increase in tundra shrubs and litter will lead to an increase in heterotrophic, biopolymer-degrading bacterial taxa and a higher incidence of potentially lichenivorous or lichenopathogenic bacteria, such as shown for the plant phyllosphere (Aydogan et al. 2018). Thus, in terms of taxonomic composition, we expected an increase in detritivorous taxa, endosymbionts and pathogens of fungi such as chitinolytic bacteria (Kobayashi and Crouch 2009), whereas the relative abundance of cold-adapted and facultatively lithotrophic bacteria may decrease.

We also hypothesized that the potentially metabolically active (16S rRNA based) community is more response to the warming treatment than the total bacterial community, as has been shown for the effect of drought (Bastida et al. 2017).

## 2 Methods

### 2.1 Study site and experimental design

The study site is located in a *Betula nana* heath in the Icelandic central highlands (65°16’N, 20°15’W) at an altidude of 450 m. According to Köppen’s climate definitions, the sampling site, called Auðkúluheiði (65°16’N, 20°15’W, 480 m above sea level) is situated in the lower Arctic. The vegetation is characterized as a relatively species-rich dwarf shrub heath, with *Betula nana* being the most dominant vascular species and the moss *Racomitrium lanuginosum* and the lichen *C. islandica* as the dominating cryptogam species (Jonsdottir et al. 2005; Arnalds 2015).

Ten open top plexiglass chambers (OTCs) were set up to simulate a warmer summer climate in August 1996 (Hollister and Webber 2000; Jonsdottir et al. 2005). The OTCs raise the mean daily temperature by 1-2 °C during summer and minimize secondary experimental effects such as differences in atmospheric gas concentration and reduction in ambient precipitation. Control plots were established adjacent to the OTCs, but without any treatment, thus exposing the environment to ambient temperatures. Per warmed (OTC) and control plot, the upper parts (2×2 cm) of five thalli were randomly selected and collected with sterile tweezers. The samples were immediately soaked in RNAlater (Ambion) to prevent RNA degradation and kept cool until storage at −80 °C. The lichen samples were collected in June 2017.

### 2.2 RNA and DNA extraction and sequencing

Prior to RNA and DNA extraction, the samples were washed with RNase free water to get rid of soil particles and RNAlater and ground for six minutes using a Mini-Beadbeater and two sterile steel beads. RNA and DNA were extracted simultaneously using the RNeasy PowerSoil Total RNA Kit (Qiagen) and the RNeasy PowerSoil DNA Elution Kit (Qiagen), following the manufacturer’s instructions. DNA and RNA concentrations were measured with a Qubit Fluorometer (Life Technologies) and purity was assessed with a NanoDrop (NanoDrop Technologies) and integrity by Bioanalyzer (Agilent Technologies). cDNA was synthesized using the High-Capacity cDNA Reverse Transcription Kit (Thermofisher) following the manufacturer’s instructions and quantified on a Qubit Fluorometer (Life Technologies). 48 DNA samples (24 from each treatment) and 48 cDNA samples (24 from each treatment) were selected to be sequenced based on the RNA and DNA quantity and quality. Library preparation and sequencing of the V3-V4 region of the 16S rRNA gene on an Illumina MiSeq platform was performed by Macrogen, Seoul, using the standard Illumina protocol and 16S rRNA gene V3-V4 primers.

### 2.3 Sequence processing

In order to obtain high-resolution data, we processed the raw sequences using the DADA2 pipeline (Callahan et al. 2016; Callahan, McMurdie, and Holmes 2017). Hereby sequences are not clustered into operational taxonomic units (OTUs), but exact sequences or amplicon sequence variants (ASVs). Forward reads were truncated at 260 bp and reverse reads at 250 bp. Assembled ASVs were assigned taxonomy using the Ribosomal Database Project (RDP) naïve Bayesian classifier (Wang et al. 2007) in DADA2 and the SILVA_132 database (Quast et al. 2013). We discarded samples with less than 10.000 non-chimeric sequences and we removed ASVs assigned to chloroplasts and mitochondria, singletons, doubletons and ASVs occurring in only 1 sample. In total, for 82 samples, 1 954 ASVs remained. The data were normalised using cumulative-sum scaling (CSS) (Paulson et al. 2013) to account for uneven sequencing depths.

The 16S rDNA based community is hereafter sometimes referred to as the DNA based community and the 16S rRNA (cDNA) based community is hereafter referred to as the cDNA based community. We interpret the cDNA based community as the ‘potentially metabolically active bacterial community’, acknowledging that 16S rRNA is not a direct indicator of activity but rather protein synthesis potential (Blazewicz et al. 2013).

### 2.4 Quantitative real-time PCR of nifH and 16S rRNA genes

We used all DNA extractions (50 replicates per treatment) for quantification of *nifH* and 16S rRNA genes, which was performed by quantitative PCR (Corbett Rotor-Gene) using the primer set PolF/PolR and 341F/534R respectively (Poly, Monrozier, and Bally 2001). The specificity of the *nifH* primers for our samples was confirmed by SANGER sequencing of 10 clone fragments. Standards for *nifH* reactions were obtained by amplifying one cloned *nifH* sequence with flanking regions of the plasmid vector (TOPO TA cloning Kit, Invitrogen). Standard curves were obtained by serial dilutions (E = 0.9 – 1.1, R^2^ = > 0.99 for all reactions). Each reaction had a volume of 20 μL, containing 1x QuantiFast SYBR Green PCR Master Mix (Qiagen), 0.2 μL of each primer (10 μM), 0.8 μL BSA (5 μg/μL), 6.8 μL RNase free water and 2 μL template. The cycling program was 5 min at 95 °C, 30 cycles of 10 s at 95 °C and 30 s at 60 °C.

### 2.5 Statistical analysis

Statistical analyses were conducted in R version 3.6.1. Richness (number of ASVs) and Shannon diversity were calculated with the R packages ‘vegan’ (Oksanen et al. 2013) and ‘phyloseq’ (McMurdie and Holmes 2013).

Differences in 16S rRNA and *nifH* gene abundance, ASV richness and Shannon diversity between the treatments and the DNA and cDNA were assessed with Bayesian (Markov chain Monte Carlo) generalised linear models using the R package ‘MCMCglmm’(Hadfield 2010) with treatment as a fixed and plot as a random factor to take into account the variation caused by pseudoreplication. We considered differences significant if the modelled 95% confidence intervals did not overlap.

For the comparisons of relative abundances between the warmed and the control samples, pseudoreplicates were averaged by the OTC or control plot they originated from. These average relative abundances were then compared using Wilcoxon rank-sum tests.

Distances between the community composition of the control and OTC samples were based on Bray-Curtis distances. The warming effect of the OTCs on the bacterial community composition was tested by permutational-MANOVA (PERMANOVA) (Anderson 2001) analysis of the Bray-Curtis distance matrices using the *adonis* function in the R package ‘vegan’ with plot as strata. Principal coordinate analysis was used to ordinate the Bray-Curtis distance matrices and to visualise the relationships between samples from OTC and control plots.

We used two methods to determine taxa sensitive to warming. First, differential abundance of bacterial genera between warmed and control samples was assessed using DESeq2 (Love, Huber, and Anders 2014) on the non-CSS normalised datasets with the R package ‘DESeq2’ (Love, Huber, and Anders 2014). The adjusted *p*-value cut-off was 0.1 (Love, Huber, and Anders 2014). Differential abundance analysis only uses ASVs present in both the OTC and control samples. The second method we used to find taxa sensitive to warming, was indicator species analysis. To find bacterial taxa indicative for the warming or the control treatment, correlation-based indicator species analysis was done with all possible site combinations using the function *multipatt* of the R package ‘indicSpecies’ (De Caceres and Legendre 2009) based on 10^3^ permutations. The indicator species analysis takes into account ASVs present in both OTC and control samples, but also ASVs present in only one of the treatments. We combined results of the DESeq2 and indicator species analysis for a final list of ASVs sensitive to warming.

## 3 Results

### 3.1 Effect of OTC treatment on ASV richness, diversity and community structure

The ASV richness and Shannon diversity of the bacterial communities associated with *C. islandica* were not significantly affected by the warming treatment (Figure 1, Supplementary Table 3). A significant difference was found for the richness, which was higher for the cDNA-based bacterial community than the DNA-based community in the warmed treatment (MCMCglmm, P = 0.024). The cDNA-based Shannon index tended to be higher than the DNA-based Shannon index (MCMCglmm, P = 0.076) (Figure 1, Supplementary Table 3).

**Figure 1.**
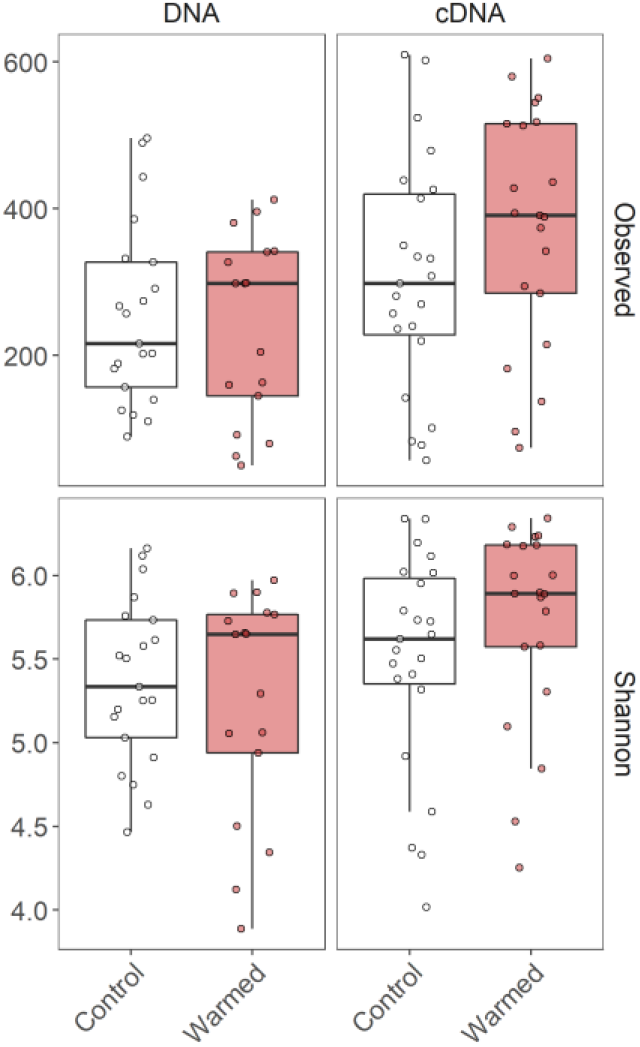
ASV richness (‘Observed’ in the figure) and Shannon diversity the DNA and cDNA-based bacterial communities associated with the lichen *Cetraria islandica* in control and warmed samples. Values for all replicates are shown. MCMC generalised linear models were used to test for differences between the treatments and the DNA and cDNA, but no differences were found between the treatments. Note that the ASV richness (‘Observed’) was higher for the cDNA-than in the DNA-based bacterial communities in the warmed treatment (pMCMC = 0.024) and the Shannon index tended to be higher for the cDNA-than the DNA-based community in the warmed treatment (pMCMC = 0.076).

Some level of clustering between the control and warmed samples could be observed in the principal coordinates analysis (Figure 3). Based on the results of a PERMANOVA, the warmed lichen associated bacterial communities were significantly different from the communities in the control plots (DNA: R^2^ = 0.07 and P = 0.001; RNA: R^2^ = 0.07 and P = 0.005) (Supplementary Table 3).

### 3.2 Effect of OTC treatment on the taxonomic composition of the *C. islandica* bacteriota

The bacterial community found associated with the lichen *C. islandica* is described at the phylum level (Figure 3A) and at the class and order level (Figure 4). No clear differences were found for the relative abundance at the phylum level between the control and warmed treatment for the total bacteria community (Figure 3A). Similarly, we did not detected differences for the potentially active bacterial community at the phylum level (Figure 3A), nor in 16S rRNA gene abundance (Figure 3B). The total bacterial community was dominated by Proteobacteria and Acidobacteria (DNA: 58% and 34% average relative abundance across all control and warmed samples, for Proteobacteria and Acidobacteria, respectively). Proteobacteria and Acidobacteria were also the main potentially active phyla in our samples (cDNA: 63% and 29%, respectively) (Figure 3A). At lower taxonomic level, the orders Acetobacterales and Acidobacteriales were the dominant taxa (DNA: 44%; cDNA: 51%, DNA: 34%; cDNA 29%, Figure 4A). Within the acidobacterial family Acetobacteraceae, about 14% could not be assigned to a genus (Supplementary Figure 2). The most abundant and potentially active genera were the proteobacterial genera *Acidiphilium* (DNA: 8%, cDNA: 11%) and *Endobacter* (DNA: 19%, cDNA: 20%) and the acidobacterial genera *Bryocella* (DNA: 10%, cDNA: 9%) and *Granulicella* (DNA: 15%, cDNA: 11%) (Supplementary Figure 2).

**Figure 2.**
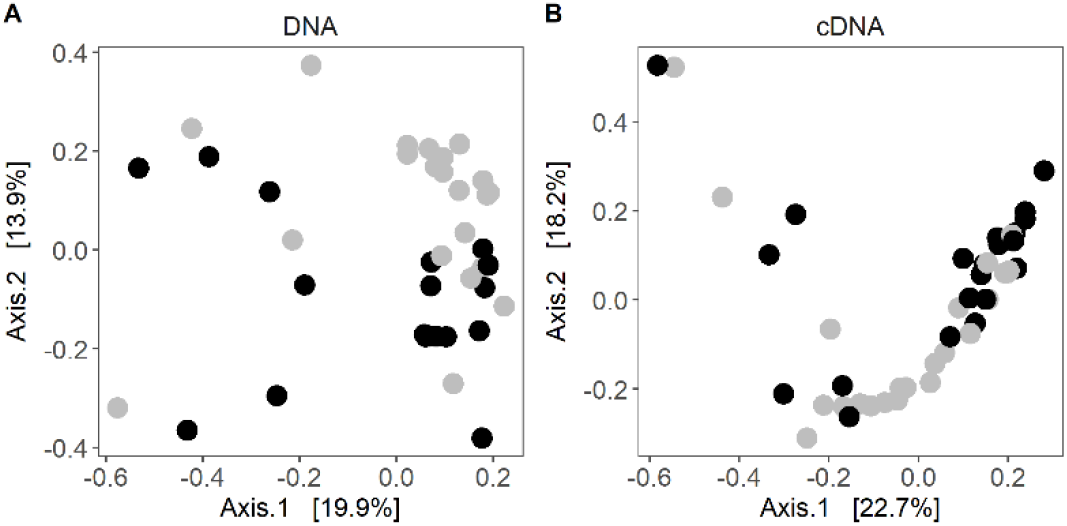
Principal coordinate analysis of bray-curtis distances of the (A) DNA-based community composition and (B) cDNA-based community composition of the lichen *Cetraria islandica*. Warmed samples are represented as black circles and control samples as white grey circles. PERMANOVA showed significant differentiation between control and warmed communities (DNA: R^2^ = 0.07 and P = 0.001; cDNA: R^2^ = 0.07 and P = 0.005).

295 ASVs were only detected in the cDNA-based samples and not in the DNA-based samples. These taxa belonged to abundant genera such as *Endobacter* and *Acidiphilium* (Supplementary Table 1 and 2). Genera that were exclusively found in the cDNA-based community were *Lactococcus*, *Lachnospiraceae NK4A136 group*, *Ktedonobacter*, *Methylovirgula*, *Frigoribacterium*, *Amnibacterium*, *Rhizobacter*, *Telmatobacter* and *Kineosporia Acidiphilium* (Supplementary Table 1 and 2).

The bacterial load was on average 671 and 1944 16S rRNA copies per ng DNA for control and warmed plots, respectively (Figure 3B). However, no differences could be found between the overall 16S rRNA gene copy numbers, nor *nifH* gene copy numbers in the control and warmed plots (Figure 3B, Supplementary Figure 1, Supplementary Table 3).

**Figure 3.**
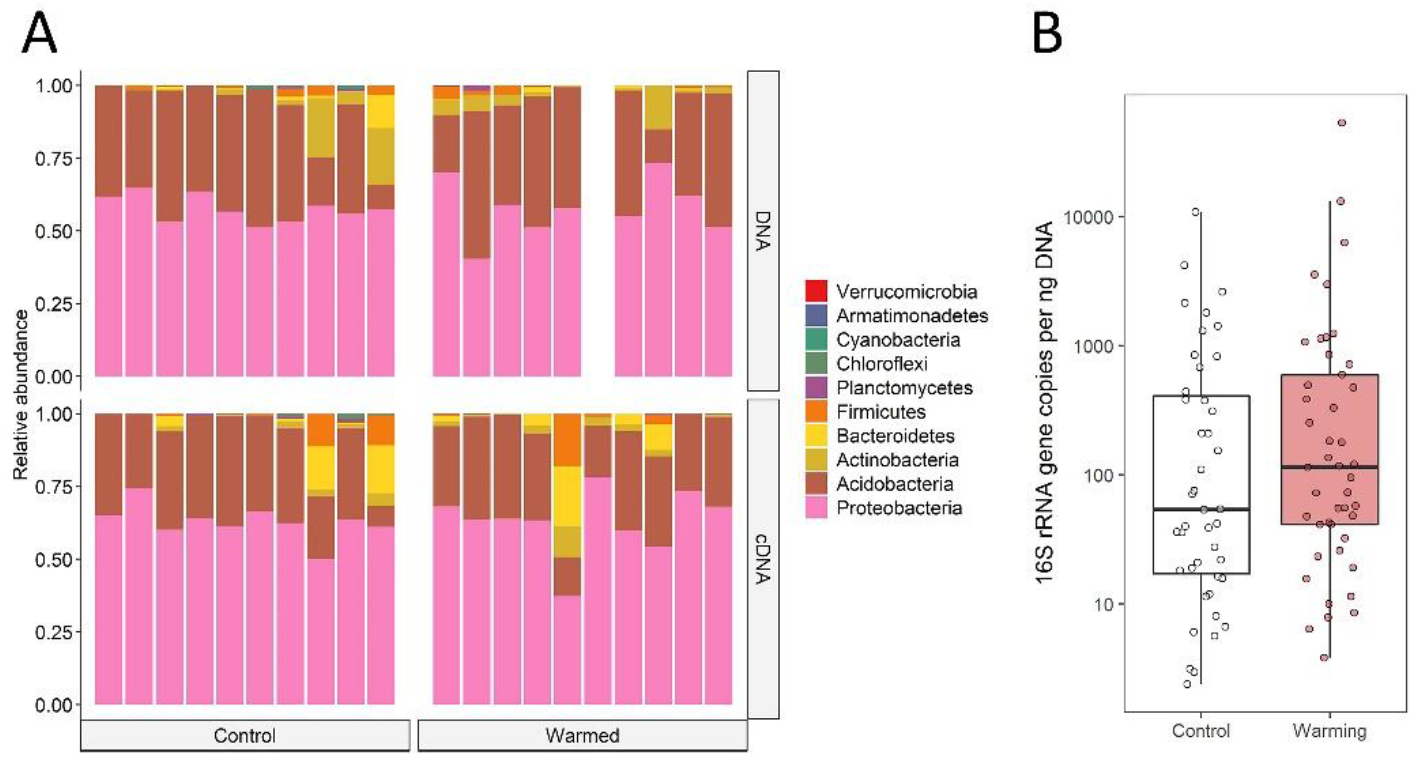
A) Barplots of the bacterial community composition found associated with the lichen C. islandica in the control and warmed plots. Data are presented at the phylum level for DNA and cDNA representing relative abundance and potential metabolically active taxa, respectively. For one of the DNA samples of the warming treatment, the number of non-chimeric sequences was less than 10.000 reads, thus it was discarded. B) 16S rRNA gene abundance per ng extracted DNA in control (white) and warmed (red) samples. All replicates are shown and differences in gene abundance between the treatments were tested using MCMC generalised linear models, but no differences could be detected.

In the DNA-based samples, the only proteobacterial class found significantly affected by warming were the Gammaproteobacteria with an increase of 50% in the warmed samples (Wilcoxon rank-sum test, *P* = 0.033; Figure 4A). Most of this increase was due to an increase in relative abundance of the order Betaproteobacteriales (*P* = 0.046) (Figure 4B). The order Sphingomonadales (Alphaproteobacteria) increased by a factor of 2.5 in relative abundance (*P* = 0.009) (Figure 4B). In the potentially active bacterial community, we did not deteted differences in relative abundance on Proteobacterial class level between the control and warmed treatment (Figure 4A). Changes were found on order level with the alphaproteobacterial order Caulobacterales being twice as potentially active in the warmed plots (*P* = 0.038) than in the control. Similarly, the order Sphingomonadales was 2.7 times more active in the warmed treatment (*P* = 0.007) (Figure 4B). The order Diplorickettsiales (Gammaproteobacteria) was four times as potentially active in the warmed plots (*P* = 0.040) (Figure 4B).

**Figure 4.**
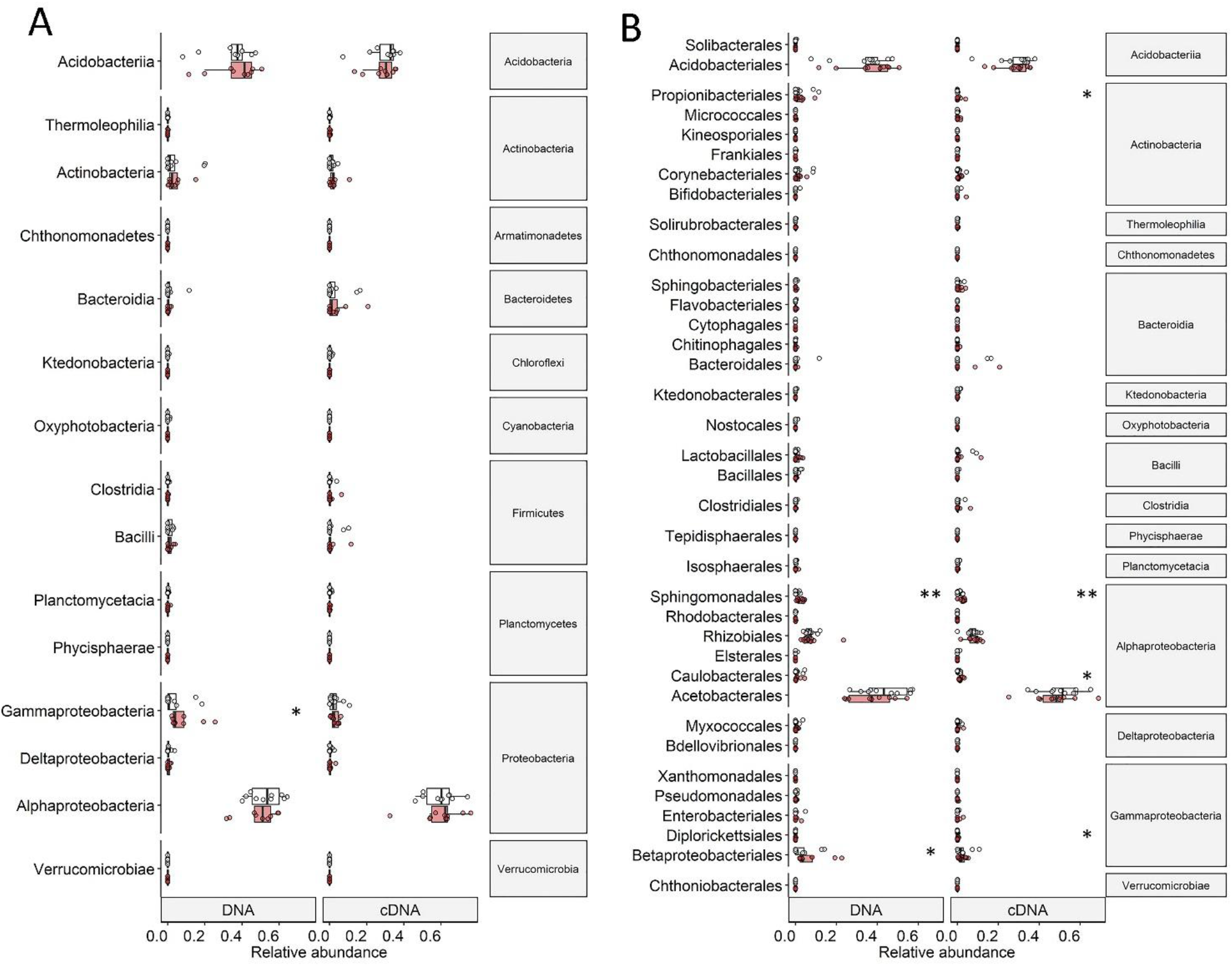
Relative abundances of A) classes and B) orders of DNA- and cDNA-based bacterial communities associated with the lichen Cetraria islandica in control (white) and warmed (red) samples. Points indicate average relative abundance values per control or warmed plot. Boxplots represent minimum values, first quartiles, medians, third quartiles and maximum values. Significance levels (* < 0.05, ** < 0.01) are based on Wilcoxon rank-sum tests.

While the phylum Actinobacteria was not among the most common phyla (DNA: XX%, cDNA: YY%), its order Propionibacteriales was 20 times as potentially active in the warmed plots compared to the control plots (*P* = 0.040) (Figure 4B).

### 3.3 Treatment effect on the relative abundance of bacterial ASVs

For the DNA-based communities, we detected 61 ASVs with a higher relative abundance in the warmed samples with a total relative abundance of 1% (Supplementary Table 1). We detected 96 ASVs with a lower relative abundance in the warmed samples compared to the control samples making up 1.7% of the total abundance (Supplementary Table 1). For the cDNA-based potentially metabolic active communities, we detected 190 ASVs with a higher relative abundance in the warmed samples (2.12%) and 77 ASVs with a lower relative abundance in the warmed samples compared to the control samples (0.9%) (Supplementary Table 2).

Of the ASVs only detected in cDNA- or potentially metabolically active bacterial community, 14 ASVs had a higher relative abundance in the warmed plots. All these rare ASVs belonged to the Proteobacteria, except one ASV that was classified as Bacteroidetes.

ASVs within the Proteobacteria showed mainly increased relative abundance in the warmed samples (Figure 5, Supplementary Table 1 and Supplementary Table 2). Only ASVs classified under the genus *Acidiphilium* had more often a lower relative abundance in the warmed samples as well as a few ASVs of the genera *Acidisphaera* and *Endobacter*. In the cDNA-based samples, more proteobacterial ASVs with increased relative abundances under warming were detected than in the DNA-based samples.

**Figure 5.**
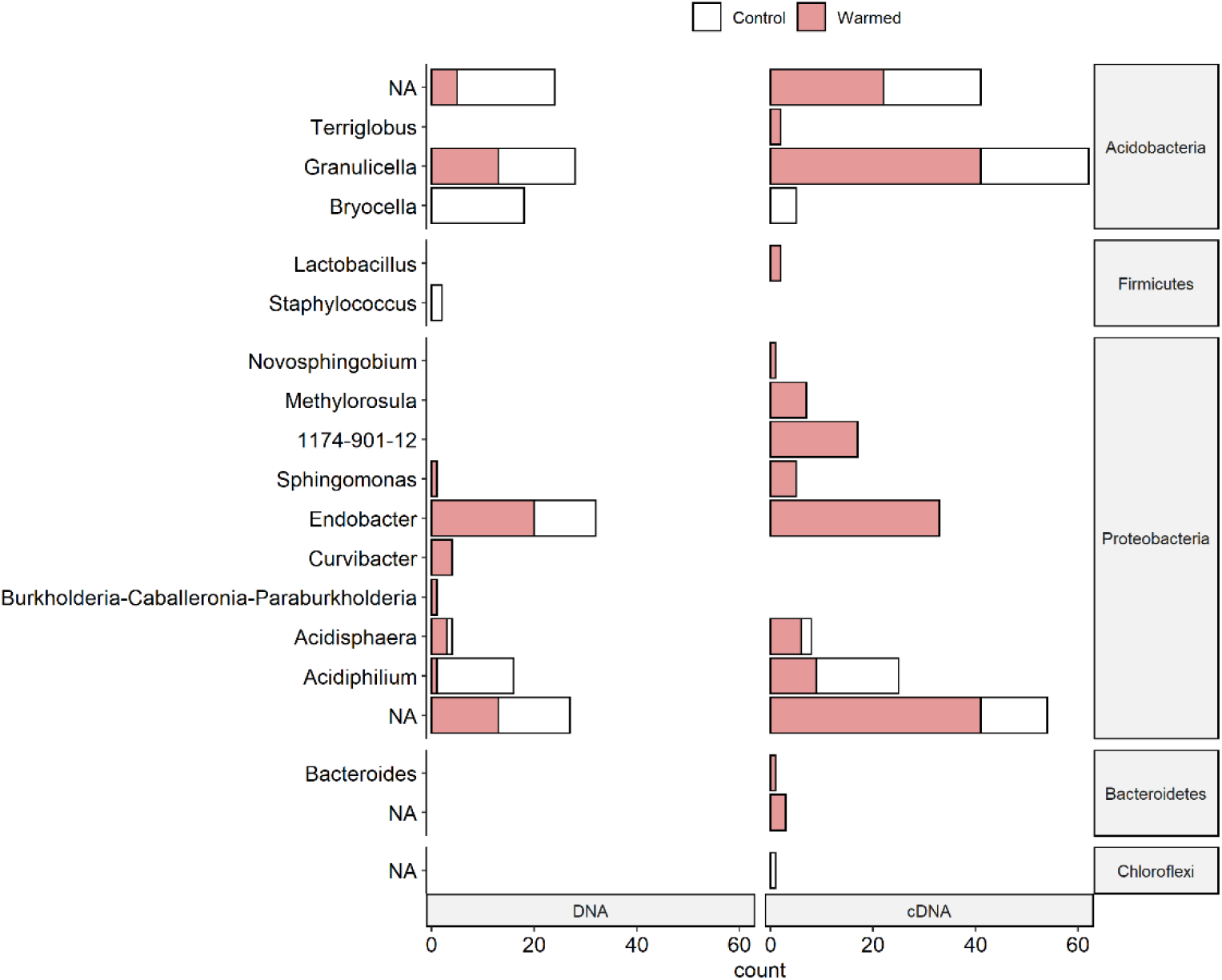
Number of ASVs (amplicon sequence variants) per genus sensitive to warming for DNA and cDNA-based bacterial communities associated with the lichen C. islandica. Sensitivity to warming was determined by differential abundance analysis (DESeq2) and indicator species analysis. ASVs not assigned to genus level are called ‘NA’. The number of ASVs indicative for the OTC (warmed) treatment is indicated in red and the number of ASVs indicative for the control treatment is indicated in white.

Acidobacterial ASVs showed mixed differences between the control and warmed samples (Figure 5, Supplementary Table 1 and Supplementary Table 2). ASVs of the genus *Bryocella* were less abundant under warming in both the DNA- and cDNA-based samples. ASVs of the genus *Granulicella* were equally more and less abundant in the warmed DNA-based samples, but had more often higher relative abundances in the warmed cDNA-based samples (Figure 5, Supplementary Table 1 and Supplementary Table 2).

## 4 Discussion

We assessed the effect of long-term (20 years) warming by open top chambers (OTCs) on the bacterial community composition associated with the lichen *C. islandica* in an Icelandic sub-Arctic alpine dwarf-shrub heath. The community was dominated by Acidobacteria and Proteobacteria <u>in both</u> total and potentially active bacterial communities in both control and warmed plots. Warming did not induce compositional or structural changes at higher taxonomical levels. Nevertheless, we found indications of multiple warming-induced shifts in the community composition at the class, order and ASV levels. The most prominent increases in relative abundance were found in several genera belonging to the Proteobacteria. Our results illustrate that the long-term warming affects the bacterial community composition of the lichen symbiosis at a fine taxonomical levels.

### 4.1 The bacterial community of *Cetraria islandica*

While the dominance of the class Alphaproteobacteria has been described as a general characteristic of lichen bacterial communities (Printzen et al. 2012; Cardinale et al. 2008; Bates et al. 2011), a striking feature of the *C. islandica* microbiome is the strong dominance of the family Acetobacteriaceae (Alphaproteobacteria). While the presence of Acetobacteriaceae in lichens has been observed before, notably in the reindeer lichen *Cladonia arbuscula* (Cardinale et al. 2008), this strong dominance does seem unusual. Indeed, the Rhizobiaceae, often held to be the dominant Alphaproteobacteria in lichens (Bates et al. 2011; Hodkinson et al. 2012), only make up a minor part of the *C. islandica* bacteriome. The second dominant group in the *C. islandica* bacteriome are members of the family Acidobacteriaceae (Acidobacteria). Even at the genus level, the *C. islandica* bacteriome is surprisingly homogeneous, with approximately half of the reads being assigned to only four genera, the acetobacterial genera *Endobacter* and *Acidiphilium*, and the acidobacterial genera *Granulicella* and *Bryocella*. This pronounced dominance of presumptively acidophilic taxa is noteworthy. Acidobacteria were also reported earlier in living parts of bog and tundra (Pankratov 2012) and lichens in Alpine soil crusts (Muggia et al. 2013). The Acidobacteria might obtain their C source from *C. islandica* by the degradation of oligosaccharides and chitin, the major component of fungal cell walls (Lladó et al. 2016).

Lichen bacterial communities are specific to certain parts of the lichen. For instance, it has been suggested that older parts of lichens harbour increased amounts of Acidobacteria (Grube et al. 2012). Mat-forming lichens such as cetraroid species are physiologically active in the apices, while the bases are composed of dead or senescing mycobiont (Grube 2010; Crittenden 1991). As we only sampled from the uppermost thalli where growth takes place, the presence of acidophilic bacteria is more likely explained by organic acid secondary metabolites produced by *C. islandica*, such as protolichesterinic and fumaroprotocetraric acids (Xu et al. 2018).

Another feature of the *C. islandica* microbiota is the difference between the cDNA (potentially metabolically active) and DNA-based community. The richness of the cDNA-based communities was higher compared the DNA-based communities. One possible explanation for this is that the detection of taxa in 16S rRNA sequences, but not in 16S rDNA sequences, can occur when rare taxa have a high metabolic potential. The occurrence of these ‘phantom taxa’ could be that as a result of the cDNA synthesis errors that do not occur in the rDNA samples are introduced in the rRNA sequences. Another explanation could be variation in metabolic activity among taxa (Klein et al. 2016; Campbell et al. 2011; Baldrian et al. 2012; Jia et al. 2019). Rare taxa have been observed to be disproportionally active compared to abundant members (Jones and Lennon 2010) and thereby might contribute more to ecosystem functioning than one would expect based on their abundance (Jousset et al. 2017). The rare biosphere of *C. islandica* was mostly composed of not assigned genera and members of the genera *Endobacter*, *Acidiphilium*, *Lactococcus*, *Mucilaginibacter* and *Bacteroides*. The fermenting bacteria from the genus *Lactococcus* have been described before in a bioreactor as being rare while having high potential activity levels (Lawson et al. 2015). The presence of rare microbes might for instance speed up the degradation of recalcitrant organic matter, such as chitin in the lichen (Peter et al. 2011).

As *C. islandica* is a chlorolichen and does not have N_2_-fixing Cyanobacteria as a photobiont, this raises the question if other taxa could be N_2_-fixers. The presence of *nifH* genes indicates that potential N_2_-fixers are present in the *C. islandica* bacterial community. Indeed we detected putative N_2_-fixers such as *Curvibacter* (Ding 2004) and members of the Burkholderiaceae. On the other hand, lichens might also obtain nutrients dissolved in precipitation or through runoff from taller vegetation, or via a moisture gradient resulting in upward movement of soil moisture and dissolved nutrients (Longton 1992). Therefore, the associated microbiota of *C. islandica* might not play such a big role in N_2_-fixation.

The *C. islandica*-associated microbiota was found to be markedly different to that of the moss *Racomitrium lanuginosum* which was studied in the same warming experiment (Klarenberg et al. 2019), further supporting the host-specific selection of bacteria from the environment and symbiotic nature of both bryophyte and lichen holobionts proposed in the recent literature (Aschenbrenner et al. 2016; Holland-Moritz et al. 2018). Specifically, we found that *C. islandica* harboured a less rich and diverse bacterial community than *R. lanuginosum*, and the microbiota composition was profoundly different. Whereas the moss was dominated by the genera *Haliangium*, *Acidiphilium*, *Nostoc*, *Conexibacter*, *Granulicella*, *Solibacter* and *Bryobacter*, the lichen was dominated by a few genera (*Bryocella*, *Granulicella*, *Acidiphilium* and *Endobacter*) as reported herein. The same difference between bacterial diversity of a lichen and a moss was shown by Aschenbrenner et al. (2017).

### 4.2 The effect of OTC warming on bacterial richness, diversity and community structure

While we did not see any significant changes in richness of diversity of the bacterial community with warming, the warmed bacterial community structure significantly differed from the control community, both for the total as well as the potentially active communities. Overall, the potentially active community tended to be more affected by warming than the total bacterial community. For instance, more indicator taxa were found in the potentially active community and many more of these indicators were found in the warmed treatment.

At a coarse taxonomic level, the bacterial community structure was quite similar between the control and warmed treatment. One possible explanation for the similarity between the richness, diversity and composition of the warmed and control lichen bacterial communities could be that over long periods of warming bacterial communities acclimatise (Bradford et al. 2008; Crowther and Bradford 2013; Romero-Olivares, Allison, and Treseder 2017). Nonetheless, at lower taxonomic levels (class, order and ASV) we detected differences in relative abundances. Shifts in individual taxa can affect microbe-microbe and microbe-host interactions and potentially change functionality or stability of the lichen-associated bacterial communities and thereby influence host health and ecosystem functioning, as proposed for plant-microbiomes (Aydogan et al. 2018; van der Heijden and Hartmann 2016; Agler et al. 2016).

Long-term warming decreased the relative abundance of ASVs belonging to the Acidobacterial genera *Granulicella* and *Bryocella* and the alphaproteobacterial genus *Acidiphilium* in the total bacterial communities. *Acidiphilium* and *Granulicella* have been observed in other lichen microbiomes (Pankratov 2012; Park et al. 2016; Bates et al. 2011; Aschenbrenner et al. 2017). These genera are chemoorganotrophic or chemolitotrophic and might thus survive on C sources present in the lichen thallus. *Granulicella* encompasses several acidophilic, cold-adapted species described from tundra soil isolates (Mannisto et al. 2012). It has hydrolytic properties such as the ability to degrade chitin (Pankratov and Dedysh 2010; Pankratov 2012; Park et al. 2016; Belova et al. 2018), which suggest a role for these bacteria in the degradation of senescing lichen thalli. While some ASVs of the genus *Granulicella* decreased in relative abundance, the increased potential activity might result in increased degradation of dead lichen material. In contrast, the decreased relative abundance and potential activity of *Bryocella* and *Acidiphilium* might result in slower degradation of dead lichen material. Strikingly, the genus *Acidiphilium* showed an increase in relative abundance in a moss microbiome in the same warming experiment (Klarenberg et al. 2019). This suggests that the response of the lichen and moss microbiomes to environmental change are species specific and could lead to different outcomes for the moss and the lichen.

The alphaproteobacterial genera *Acidisphaera*, *Sphingomonas and Endobacter* showed an increased relative abundance and potential activity with warming. *Sphingomonas* and *Acidisphaera* have been identified in other lichen bacterial communities (Cardinale et al. 2008). *Endobacter* is a poorly characterized genus of which only one species has been described (Ramirez-Bahena et al. 2013). *Acidisphaera* is chemoorganotrophic and contains bacteriochlorophyll (Hiraishi et al. 2000). *Sphingomonas* is known for its ability to degrade plant biomass, the utilisation of recalcitrant matter in oligotrophic environments, and the use of sulfonated compounds as sources of C and sulfur (Aylward et al. 2013). The increase in relative abundance and potential activity of these genera in the warmed conditions might enhance C and nutrient availability in the lichen thalli.

Overall, all genera that dominated the bacterial community of *C. islandica* contained ASVs that were affected by the warming treatment in their relative abundances and potential metabolic activity. The genera that were affected in relative abundance are likely to play roles in nutrient recycling and supply in the lichen symbiosis. As most of these ASVs increased in relative abundance with warming, nutrient turnover in the lichen might be accelerated.

Temperature-induced changes in the amount of litter, surrounding plant species composition, lichen traits such as thallus nutrient content, soil organic matter content, and soil moisture are environmental factors that could potentially influence the lichen bacterial community composition (Vandenkoornhuyse et al. 2015; Koyama et al. 2018; Sayer et al. 2017). Warming could also affect the secondary metabolites of the lichen (Asplund, Siegenthaler, and Gauslaa 2017) and thereby alter the composition of the lichen microbiota. Therefore, our results suggest that a warmer tundra can lead to changes in lichen bacterial communities at a fine taxonomic level. The lichen microbiome plays an important role in the growth of lichens and climate-driven changes in the lichen microbiota, irrespective of whether they are due to direct or indirect effects of climate change, might thus affect decomposition of lichens and thereby nutrient cycling in sub-Arctic ecosystems.

## 6 Author contributions

IJ, IK and OV designed the study. IK conducted the sampling, the laboratory work, the bioinformatics processing and the statistical analysis. CK performed the qPCR measurements. IK wrote the manuscript with contributions from OV, IJ, CK and DW.

## 8 Acknowledgements

We kindly thank Margrét Auður Sigurbjörnsdóttir for providing insightful comments on the manuscript.

## References

Agler, Matthew T., Jonas Ruhe, Samuel Kroll, Constanze Morhenn, Sang-Tae Kim, Detlef Weigel, and Eric M. Kemen. 2016. “Microbial Hub Taxa Link Host and Abiotic Factors to Plant Microbiome Variation.” Edited by Matthew K. Waldor. PLOS Biology 14 (1): e1002352. https://doi.org/10.1371/journal.pbio.1002352.

Alatalo, Juha M., Annika K. Jägerbrand, Shengbin Chen, and Ulf Molau. 2017. “Responses of Lichen Communities to 18 Years of Natural and Experimental Warming.” Annals of Botany 120 (1): 159–70. https://doi.org/10.1093/aob/mcx053.

Almendras, Katerin, Jaime García, Margarita Carú, and Julieta Orlando. 2018. “Nitrogen-Fixing Bacteria Associated with Peltigera Cyanolichens and Cladonia Chlorolichens.” Molecules 23 (12): 3077. https://doi.org/10.3390/molecules23123077.

Anderson, Marti J. 2001. “A New Method for Non-Parametric Multivariate Analysis of Variance.” Austral Ecology 26 (1): 32–46.

Arnalds, Olafur. 2015. The Soils of Iceland. World Soils Book Series. Dordrecht: Springer Netherlands. http://link.springer.com/10.1007/978-94-017-9621-7.

Aschenbrenner, Ines Aline, Tomislav Cernava, Gabriele Berg, and Martin Grube. 2016. “Understanding Microbial Multi-Species Symbioses.” Frontiers in Microbiology 7 (February). https://doi.org/10.3389/fmicb.2016.00180.

Aschenbrenner, Ines Aline, Tomislav Cernava, Armin Erlacher, Gabriele Berg, and Martin Grube. 2017. “Differential Sharing and Distinct Co-Occurrence Networks among Spatially Close Bacterial Microbiota of Bark, Mosses and Lichens.” Molecular Ecology 26 (10): 2826–38. https://doi.org/10.1111/mec.14070.

Asplund, Johan, Andy Siegenthaler, and Yngvar Gauslaa. 2017. “Simulated Global Warming Increases Usnic Acid but Reduces Perlatolic Acid in the Mat-Forming Terricolous Lichen *Cladonia Stellaris*.” The Lichenologist 49 (3): 269–74. https://doi.org/10.1017/S0024282917000159.

Asplund, Johan, and David A. Wardle. 2017. “How Lichens Impact on Terrestrial Community and Ecosystem Properties.” Biological Reviews 92 (3): 1720–38. https://doi.org/10.1111/brv.12305.

Aydogan, Ebru L., Gerald Moser, Christoph Müller, Peter Kämpfer, and Stefanie P. Glaeser. 2018. “Long-Term Warming Shifts the Composition of Bacterial Communities in the Phyllosphere of Galium Album in a Permanent Grassland Field-Experiment.” Frontiers in Microbiology 9 (144). https://doi.org/10.3389/fmicb.2018.00144.

Aylward, Frank O., Bradon R. McDonald, Sandra M. Adams, Alejandra Valenzuela, Rebeccah A. Schmidt, Lynne A. Goodwin, Tanja Woyke, Cameron R. Currie, Garret Suen, and Michael Poulsen. 2013. “Comparison of 26 Sphingomonad Genomes Reveals Diverse Environmental Adaptations and Biodegradative Capabilities.” Applied and Environmental Microbiology 79 (12): 3724–33. https://doi.org/10.1128/AEM.00518-13.

Baldrian, Petr, Miroslav Kolařík, Martina Štursová, Jan Kopecký, Vendula Valášková, Tomáš Větrovský, Lucia Žifčáková, et al. 2012. “Active and Total Microbial Communities in Forest Soil Are Largely Different and Highly Stratified during Decomposition.” The ISME Journal 6 (2): 248–58. https://doi.org/10.1038/ismej.2011.95.

Bastida, Felipe, Irene F. Torres, Manuela Andrés-Abellán, Petr Baldrian, Rubén López-Mondéjar, Tomáš Větrovský, Hans H. Richnow, et al. 2017. “Differential Sensitivity of Total and Active Soil Microbial Communities to Drought and Forest Management.” Global Change Biology 23 (10): 4185–4203. https://doi.org/10.1111/gcb.13790.

Bates, S. T., G. W. G. Cropsey, J. G. Caporaso, R. Knight, and N. Fierer. 2011. “Bacterial Communities Associated with the Lichen Symbiosis.” Applied and Environmental Microbiology 77 (4): 1309–14. https://doi.org/10.1128/AEM.02257-10.

Belova, Svetlana E., Nikolai V. Ravin, Timofey A. Pankratov, Andrey L. Rakitin, Anastasia A. Ivanova, Alexey V. Beletsky, Andrey V. Mardanov, Jaap S. Sinninghe Damsté, and Svetlana N. Dedysh. 2018. “Hydrolytic Capabilities as a Key to Environmental Success: Chitinolytic and Cellulolytic Acidobacteria From Acidic Sub-Arctic Soils and Boreal Peatlands.” Frontiers in Microbiology 9. https://doi.org/10.3389/fmicb.2018.02775.

Biasi, Christina, Hildegard Meyer, Olga Rusalimova, Rainer Hämmerle, Christina Kaiser, Christian Baranyi, Holger Daims, Nikolaj Lashchinsky, Pavel Barsukov, and Andreas Richter. 2008. “Initial Effects of Experimental Warming on Carbon Exchange Rates, Plant Growth and Microbial Dynamics of a Lichen-Rich Dwarf Shrub Tundra in Siberia.” Plant and Soil 307 (1–2): 191–205. https://doi.org/10.1007/s11104-008-9596-2.

Bjelland, Torbjørg, Martin Grube, Solveig Hoem, Steffen L. Jorgensen, Frida Lise Daae, Ingunn H. Thorseth, and Lise Øvreås. 2011. “Microbial Metacommunities in the Lichen-Rock Habitat: Microbial Metacommunities in the Lichen-Rock Habitat.” Environmental Microbiology Reports 3 (4): 434–42. https://doi.org/10.1111/j.1758-2229.2010.00206.x.

Blazewicz, Steven J., Romain L. Barnard, Rebecca A. Daly, and Mary K. Firestone. 2013. “Evaluating RRNA as an Indicator of Microbial Activity in Environmental Communities: Limitations and Uses.” The ISME Journal 7: 2061–68.

Bradford, Mark A., Christian A. Davies, Serita D. Frey, Thomas R. Maddox, Jerry M. Melillo, Jacqueline E. Mohan, James F. Reynolds, Kathleen K. Treseder, and Matthew D. Wallenstein. 2008. “Thermal Adaptation of Soil Microbial Respiration to Elevated Temperature.” Ecology Letters 11 (12): 1316–27. https://doi.org/10.1111/j.1461-0248.2008.01251.x.

Callahan, Benjamin J., Paul J. McMurdie, and Susan P. Holmes. 2017. “Exact Sequence Variants Should Replace Operational Taxonomic Units in Marker-Gene Data Analysis.” The ISME Journal 11 (December): 2639–43. https://doi.org/10.1038/ismej.2017.119.

Callahan, Benjamin J., Paul J. McMurdie, Michael J. Rosen, Andrew W. Han, Amy Jo A. Johnson, and Susan P. Holmes. 2016. “DADA2: High-Resolution Sample Inference from Illumina Amplicon Data.” Nature Methods 13 (7): 581–83. https://doi.org/10.1038/nmeth.3869.

Campbell, Barbara J., Liying Yu, John F. Heidelberg, and David L. Kirchman. 2011. “Activity of Abundant and Rare Bacteria in a Coastal Ocean.” Proceedings of the National Academy of Sciences 108 (31): 12776–81. https://doi.org/10.1073/pnas.1101405108.

Cardinale, Massimiliano, Martin Grube, João Vieirade Castro, Henry Müller, and Gabriele Berg. 2012. “Bacterial Taxa Associated with the Lung Lichen Lobaria Pulmonaria Are Differentially Shaped by Geography and Habitat.” FEMS Microbiology Letters 329 (2): 111–15. https://doi.org/10.1111/j.1574-6968.2012.02508.x.

Cardinale, Massimiliano, Anna Maria Puglia, and Martin Grube. 2006. “Molecular Analysis of Lichen-Associated Bacterial Communities: Lichen-Associated Bacterial Communities.” FEMS Microbiology Ecology 57 (3): 484–95. https://doi.org/10.1111/j.1574-6941.2006.00133.x.

Cardinale, Massimiliano, Jana Steinová, Johannes Rabensteiner, Gabriele Berg, and Martin Grube. 2012. “Age, Sun and Substrate: Triggers of Bacterial Communities in Lichens: Ecology of Lichen-Associated Bacterial Communities.” Environmental Microbiology Reports 4 (1): 23–28. https://doi.org/10.1111/j.1758-2229.2011.00272.x.

Cardinale, Massimiliano, JoÃ£o Vieira de Castro, Henry MÃ¼ller, Gabriele Berg, and Martin Grube. 2008. “In Situ Analysis of the Bacterial Community Associated with the Reindeer Lichen Cladonia Arbuscula Reveals Predominance of Alphaproteobacteria: Lichen-Associated Bacterial Community.” FEMS Microbiology Ecology 66 (1): 63–71. https://doi.org/10.1111/j.1574-6941.2008.00546.x.

Cernava, Tomislav, Ines Aline Aschenbrenner, Jung Soh, Christoph W. Sensen, Martin Grube, and Gabriele Berg. 2019. “Plasticity of a Holobiont: Desiccation Induces Fasting-like Metabolism within the Lichen Microbiota.” The ISME Journal 13 (2): 547. https://doi.org/10.1038/s41396-018-0286-7.

Cernava, Tomislav, Qerimane Vasfiu, Armin Erlacher, Ines Aline Aschenbrenner, Kevin Francesconi, Martin Grube, and Gabriele Berg. 2018. “Adaptions of Lichen Microbiota Functioning Under Persistent Exposure to Arsenic Contamination.” Frontiers in Microbiology 9. https://doi.org/10.3389/fmicb.2018.02959.

Coleine, Claudia, Jason E. Stajich, Nuttapon Pombubpa, Laura Zucconi, Silvano Onofri, Fabiana Canini, and Laura Selbmann. 2019. “Altitude and Fungal Diversity Influence the Structure of Antarctic Cryptoendolithic Bacteria Communities.” Environmental Microbiology Reports 11 (5): 718–26. https://doi.org/10.1111/1758-2229.12788.

Cornelissen, J. H. C., T. V. Callaghan, J. M. Alatalo, A. Michelsen, E. Graglia, A. E. Hartley, D. S. Hik, et al. 2001. “Global Change and Arctic Ecosystems: Is Lichen Decline a Function of Increases in Vascular Plant Biomass?” Journal of Ecology 89 (6): 984–94. https://doi.org/10.1111/j.1365-2745.2001.00625.x.

Cornelissen, J. H. C., S. I. Lang, N. A. Soudzilovskaia, and H. J. During. 2007. “Comparative Cryptogam Ecology: A Review of Bryophyte and Lichen Traits That Drive Biogeochemistry.” Annals of Botany 99 (5): 987–1001. https://doi.org/10.1093/aob/mcm030.

Crittenden, P. D. 1991. “Ecological Significance of Necromass Production In Mat-Forming Lichens.” The Lichenologist 23 (3): 323–31. https://doi.org/10.1017/S0024282991000464.

Crowther, Thomas W., and Mark A. Bradford. 2013. “Thermal Acclimation in Widespread Heterotrophic Soil Microbes.” Ecology Letters 16 (4): 469–77. https://doi.org/10.1111/ele.12069.

De Caceres, Miquel, and Pierre Legendre. 2009. “Associations between Species and Groups of Sites: Indices and Statistical Inference.” Ecology 90 (12): 3566–74.

Ding, L. 2004. “Proposals of Curvibacter Gracilis Gen. Nov., Sp. Nov. and Herbaspirillum Putei Sp. Nov. for Bacterial Strains Isolated from Well Water and Reclassification of [Pseudomonas] Huttiensis, [Pseudomonas] Lanceolata, [Aquaspirillum] Delicatum and [Aquaspirillum] Autotrophicum as Herbaspirillum Huttiense Comb. Nov., Curvibacter Lanceolatus Comb. Nov., Curvibacter Delicatus Comb. Nov. and Herbaspirillum Autotrophicum Comb. Nov.” INTERNATIONAL JOURNAL OF SYSTEMATIC AND EVOLUTIONARY MICROBIOLOGY 54 (6): 2223–30. https://doi.org/10.1099/ijs.0.02975-0.

Domaschke, S., M. Vivas, L. G. Sancho, and C. Printzen. 2013. “Ecophysiology and Genetic Structure of Polar versus Temperate Populations of the Lichen Cetraria Aculeata.” Oecologia 173 (3): 699–709. https://doi.org/10.1007/s00442-013-2670-3.

Elmendorf, Sarah C., Gregory H. R. Henry, Robert D. Hollister, Robert G. Björk, Anne D. Bjorkman, Terry V. Callaghan, Laura Siegwart Collier, et al. 2012. “Global Assessment of Experimental Climate Warming on Tundra Vegetation: Heterogeneity over Space and Time: Warming Effects on Tundra Vegetation.” Ecology Letters 15 (2): 164–75. https://doi.org/10.1111/j.1461-0248.2011.01716.x.

Fraser, Robert H., Trevor C. Lantz, Ian Olthof, Steven V. Kokelj, and Richard A. Sims. 2014. “Warming-Induced Shrub Expansion and Lichen Decline in the Western Canadian Arctic.” Ecosystems 17 (7): 1151–68. https://doi.org/10.1007/s10021-014-9783-3.

González, Ignacio, Angel Ayuso-Sacido, Annaliesa Anderson, and Olga Genilloud. 2005. “Actinomycetes Isolated from Lichens: Evaluation of Their Diversity and Detection of Biosynthetic Gene Sequences.” FEMS Microbiology Ecology 54 (3): 401–15. https://doi.org/10.1016/j.femsec.2005.05.004.

Grube, Martin. 2010. “Die Hard: Lichens.” In Symbioses and Stress, edited by Joseph Seckbach and Martin Grube, 17:509–23. Dordrecht: Springer Netherlands. https://doi.org/10.1007/978-90-481-9449-0_26.

Grube, Martin, Massimiliano Cardinale, João Vieira de Castro, Henry Müller, and Gabriele Berg. 2009. “Species-Specific Structural and Functional Diversity of Bacterial Communities in Lichen Symbioses.” The ISME Journal 3 (9): 1105–1115.

Grube, Martin, Tomislav Cernava, Jung Soh, Stephan Fuchs, Ines Aline Aschenbrenner, Christian Lassek, Uwe Wegner, et al. 2015. “Exploring Functional Contexts of Symbiotic Sustain within Lichen-Associated Bacteria by Comparative Omics.” The ISME Journal 9 (2): 412–424.

Grube, Martin, Martina Köberl, Stefan Lackner, Christian Berg, and Gabriele Berg. 2012. “Host-Parasite Interaction and Microbiome Response: Effects of Fungal Infections on the Bacterial Community of the Alpine Lichen *Solorina Crocea*.” FEMS Microbiology Ecology 82 (2): 472–81. https://doi.org/10.1111/j.1574-6941.2012.01425.x.

Hadfield, Jarrod D. 2010. “MCMC Methods for Multi-Response Generalized Linear Mixed Models: The MCMCglmm R Package.” Journal of Statistical Software 33 (1): 1–22. https://doi.org/10.18637/jss.v033.i02.

Heijden, Marcel G. A. van der, and Martin Hartmann. 2016. “Networking in the Plant Microbiome.” PLOS Biology 14 (2): e1002378. https://doi.org/10.1371/journal.pbio.1002378.

Hiraishi, A, Y Matsuzawa, T Kanbe, and N Wakao. 2000. “Acidisphaera Rubrifaciens Gen. Nov., Sp. Nov., an Aerobic Bacteriochlorophyll-Containing Bacterium Isolated from Acidic Environments.” International Journal of Systematic and Evolutionary Microbiology, 50 (4): 1539–46. https://doi.org/10.1099/00207713-50-4-1539.

Hodkinson, Brendan P., Neil R. Gottel, Christopher W. Schadt, and François Lutzoni. 2012. “Photoautotrophic Symbiont and Geography Are Major Factors Affecting Highly Structured and Diverse Bacterial Communities in the Lichen Microbiome: Prokaryotic Communities of the Lichen Microbiome.” Environmental Microbiology 14 (1): 147–61. https://doi.org/10.1111/j.1462-2920.2011.02560.x.

Hodkinson, Brendan P., and François Lutzoni. 2009. “A Microbiotic Survey of Lichen-Associated Bacteria Reveals a New Lineage from the Rhizobiales.” Symbiosis 49 (3): 163–180.

Holland-Moritz, Hannah, Julia Stuart, Lily R. Lewis, Samantha Miller, Michelle C. Mack, Stuart F. McDaniel, and Noah Fierer. 2018. “Novel Bacterial Lineages Associated with Boreal Moss Species.” Environmental Microbiology 20 (7): 2625–38. https://doi.org/10.1111/1462-2920.14288.

Hollister, Robert D., and Patrick J. Webber. 2000. “Biotic Validation of Small Open-Top Chambers in a Tundra Ecosystem.” Global Change Biology 6 (7): 835–42. https://doi.org/10.1046/j.1365-2486.2000.00363.x.

IPCC. 2019. “Special Report on the Ocean and Cryosphere in a Changing Climate (SROCC).” https://report.ipcc.ch/srocc/pdf/SROCC_FinalDraft_FullReport.pdf.

Jia, Yangyang, Marcus H. Y. Leung, Xinzhao Tong, David Wilkins, and Patrick K. H. Lee. 2019. “Rare Taxa Exhibit Disproportionate Cell-Level Metabolic Activity in Enriched Anaerobic Digestion Microbial Communities.” MSystems 4 (1). https://doi.org/10.1128/mSystems.00208-18.

Jones, Stuart E., and Jay T. Lennon. 2010. “Dormancy Contributes to the Maintenance of Microbial Diversity.” Proceedings of the National Academy of Sciences 107 (13): 5881–86. https://doi.org/10.1073/pnas.0912765107.

Jonsdottir, Ingibjorg S., Borgthor Magnusson, Jon Gudmundsson, Asrun Elmarsdottir, and Hreinn Hjartarson. 2005. “Variable Sensitivity of Plant Communities in Iceland to Experimental Warming.” Global Change Biology 11 (4): 553–63. https://doi.org/10.1111/j.1365-2486.2005.00928.x.

Jousset, Alexandre, Christina Bienhold, Antonis Chatzinotas, Laure Gallien, Angélique Gobet, Viola Kurm, Kirsten Küsel, et al. 2017. “Where Less May Be More: How the Rare Biosphere Pulls Ecosystems Strings.” The ISME Journal 11 (4): 853–62. https://doi.org/10.1038/ismej.2016.174.

Kärnefelt, Ingvar, Jan-Eric Mattsson, and Arne Thell. 1993. “The Lichen Genera Arctocetraria, Cetraria, and Cetrariella (Parmeliaceae) and Their Presumed Evolutionary Affinities.” The Bryologist 96 (3): 394–404. https://doi.org/10.2307/3243869.

Klarenberg, Ingeborg J., Christoph Keuschnig, Ana J. Russi Colmenares, Anne D. Jungblut, Ingibjörg S. Jónsdóttir, and Oddur Vilhelmsson. 2019. “Long-Term Warming Effects on the Microbiome and Nitrogen Fixation Associated with the Moss Racomitrium Lanuginosum in a Subarctic Alpine Heathland.” BioRxiv, November, 838581. https://doi.org/10.1101/838581.

Klein, Ann M., Brendan J. M. Bohannan, Daniel A. Jaffe, David A. Levin, and Jessica L. Green. 2016. “Molecular Evidence for Metabolically Active Bacteria in the Atmosphere.” Frontiers in Microbiology 7. https://doi.org/10.3389/fmicb.2016.00772.

Kobayashi, Donald Y., and Jo Anne Crouch. 2009. “Bacterial/Fungal Interactions: From Pathogens to Mutualistic Endosymbionts.” Annual Review of Phytopathology 47 (1): 63–82. https://doi.org/10.1146/annurev-phyto-080508-081729.

Koyama, Akihiro, J. Megan Steinweg, Michelle L. Haddix, Jeffrey S. Dukes, and Matthew D. Wallenstein. 2018. “Soil Bacterial Community Responses to Altered Precipitation and Temperature Regimes in an Old Field Grassland Are Mediated by Plants.” FEMS Microbiology Ecology 94 (1): fix156. https://doi.org/10.1093/femsec/fix156.

Lawson, Christopher E., Blake J. Strachan, Niels W. Hanson, Aria S. Hahn, Eric R. Hall, Barry Rabinowitz, Donald S. Mavinic, William D. Ramey, and Steven J. Hallam. 2015. “Rare Taxa Have Potential to Make Metabolic Contributions in Enhanced Biological Phosphorus Removal Ecosystems.” Environmental Microbiology 17 (12): 4979–93. https://doi.org/10.1111/1462-2920.12875.

Lladó, Salvador, Lucia Žifčáková, Tomáš Větrovský, Ivana Eichlerová, and Petr Baldrian. 2016. “Functional Screening of Abundant Bacteria from Acidic Forest Soil Indicates the Metabolic Potential of Acidobacteria Subdivision 1 for Polysaccharide Decomposition.” Biology and Fertility of Soils 52 (2): 251–60. https://doi.org/10.1007/s00374-015-1072-6.

Longton, RE. 1992. “The Role of Bryophytes and Lichens in Terrestrial Ecosystems.” In Bryophytes and Lichens in a Changing Environment, edited by Jeffrey Bates and Farmer, AM, 32–76. Oxford: University Press. https://www.esf.edu/efb/Kimmerer/mossecology/reserve/Role_of_bryophytes_and_lichens_in_terrestrial_ecosystems.pdf.

Love, Michael I, Wolfgang Huber, and Simon Anders. 2014. “Moderated Estimation of Fold Change and Dispersion for RNA-Seq Data with DESeq2.” Genome Biology 15 (550). https://doi.org/10.1186/s13059-014-0550-8.

Mannisto, M. K., S. Rawat, V. Starovoytov, and M. M. Haggblom. 2012. “Granulicella Arctica Sp. Nov., Granulicella Mallensis Sp. Nov., Granulicella Tundricola Sp. Nov. and Granulicella Sapmiensis Sp. Nov., Novel Acidobacteria from Tundra Soil.” INTERNATIONAL JOURNAL OF SYSTEMATIC AND EVOLUTIONARY MICROBIOLOGY 62 (Pt 9): 2097–2106. https://doi.org/10.1099/ijs.0.031864-0.

McMurdie, Paul J., and Susan Holmes. 2013. “Phyloseq: An R Package for Reproducible Interactive Analysis and Graphics of Microbiome Census Data.” PLOS ONE 8 (4): e61217. https://doi.org/10.1371/journal.pone.0061217.

Muggia, Lucia, Barbara Klug, Gabriele Berg, and Martin Grube. 2013. “Localization of Bacteria in Lichens from Alpine Soil Crusts by Fluorescence in Situ Hybridization.” Applied Soil Ecology 68 (June): 20–25. https://doi.org/10.1016/j.apsoil.2013.03.008.

Mushegian, Alexandra A., Celeste N. Peterson, Christopher C. M. Baker, and Anne Pringle. 2011. “Bacterial Diversity across Individual Lichens.” Applied and Environmental Microbiology 77 (12): 4249–52. https://doi.org/10.1128/AEM.02850-10.

Myers-Smith, Isla H., Meagan M. Grabowski, Haydn J. D. Thomas, Sandra Angers-Blondin, Gergana N. Daskalova, Anne D. Bjorkman, Andrew M. Cunliffe, et al. 2019. “Eighteen Years of Ecological Monitoring Reveals Multiple Lines of Evidence for Tundra Vegetation Change.” Ecological Monographs 89 (2): e01351. https://doi.org/10.1002/ecm.1351.

Nash, Thomas H. 2008. Lichen Biology. 2nd ed. Cambridge University Press. https://doi.org/10.1017/CBO9780511790478.

Nash, Thomas H., and Astrid G. Olafsen. 1995. “Climate Change and the Ecophysiological Response of Arctic Lichens.” The Lichenologist 27 (06): 559–565.

Oksanen, Jari, F. Guillaume Blanchet, Roeland Kindt, Pierre Legendre, Peter R. Minchin, R. B. O’hara, Gavin L. Simpson, Peter Solymos, M. Henry H. Stevens, and Helene Wagner. 2013. “Package ‘Vegan.’” Community Ecology Package, Version 2 (9): 296.

Pankratov, Timofey A. 2012. “Acidobacteria in Microbial Communities of the Bog and Tundra Lichens.” Microbiology 81 (1): 51–58. https://doi.org/10.1134/S0026261711060166.

Pankratov, Timofey A., and Svetlana N. Dedysh. 2010. “Granulicella Paludicola Gen. Nov., Sp. Nov., Granulicella Pectinivorans Sp. Nov., Granulicella Aggregans Sp. Nov. and Granulicella Rosea Sp. Nov., Acidophilic, Polymer-Degrading Acidobacteria from Sphagnum Peat Bogs.” International Journal of Systematic and Evolutionary Microbiology 60 (Pt 12): 2951–59. https://doi.org/10.1099/ijs.0.021824-0.

Park, Chae Haeng, Kyung Mo Kim, Ok-Sun Kim, Gajin Jeong, and Soon Gyu Hong. 2016. “Bacterial Communities in Antarctic Lichens.” Antarctic Science 28 (06): 455–61. https://doi.org/10.1017/S0954102016000286.

Paulson, Joseph N, O Colin Stine, Héctor Corrada Bravo, and Mihai Pop. 2013. “Differential Abundance Analysis for Microbial Marker-Gene Surveys.” Nature Methods 10 (12): 1200–1202. https://doi.org/10.1038/nmeth.2658.

Peter, Hannes, Sara Beier, Stefan Bertilsson, Eva S. Lindström, Silke Langenheder, and Lars J. Tranvik. 2011. “Function-Specific Response to Depletion of Microbial Diversity.” The ISME Journal 5 (2): 351–61. https://doi.org/10.1038/ismej.2010.119.

Poly, Franck, Lucile Jocteur Monrozier, and René Bally. 2001. “Improvement in the RFLP Procedure for Studying the Diversity of NifH Genes in Communities of Nitrogen Fixers in Soil.” Research in Microbiology 152 (1): 95–103. https://doi.org/10.1016/S0923-2508(00)01172-4.

Printzen, Christian, Fernando Fernández-Mendoza, Lucia Muggia, Gabriele Berg, and Martin Grube. 2012. “Alphaproteobacterial Communities in Geographically Distant Populations of the Lichen *Cetraria Aculeata*.” FEMS Microbiology Ecology 82 (2): 316–25. https://doi.org/10.1111/j.1574-6941.2012.01358.x.

Quast, Christian, Elmar Pruesse, Pelin Yilmaz, Jan Gerken, Timmy Schweer, Pablo Yarza, Jörg Peplies, and Frank Oliver Glöckner. 2013. “The SILVA Ribosomal RNA Gene Database Project: Improved Data Processing and Web-Based Tools.” Nucleic Acids Research 41 (D1): D590–96. https://doi.org/10.1093/nar/gks1219.

Ramirez-Bahena, M. H., C. Tejedor, I. Martin, E. Velazquez, and A. Peix. 2013. “Endobacter Medicaginis Gen. Nov., Sp. Nov., Isolated from Alfalfa Nodules in an Acidic Soil.” INTERNATIONAL JOURNAL OF SYSTEMATIC AND EVOLUTIONARY MICROBIOLOGY 63 (Pt 5): 1760–65. https://doi.org/10.1099/ijs.0.041368-0.

Rolshausen, Gregor, Francesco Dal Grande, Anna D. Sadowska-Deś, Jürgen Otte, and Imke Schmitt. 2018. “Quantifying the Climatic Niche of Symbiont Partners in a Lichen Symbiosis Indicates Mutualist-Mediated Niche Expansions.” Ecography 41 (8): 1380–92. https://doi.org/10.1111/ecog.03457.

Romero-Olivares, A. L., S. D. Allison, and K. K. Treseder. 2017. “Soil Microbes and Their Response to Experimental Warming over Time: A Meta-Analysis of Field Studies.” Soil Biology and Biochemistry 107 (April): 32–40. https://doi.org/10.1016/j.soilbio.2016.12.026.

Sayer, Emma J., Anna E. Oliver, Jason D. Fridley, Andrew P. Askew, Robert T. E. Mills, and J. Philip Grime. 2017. “Links between Soil Microbial Communities and Plant Traits in a Species-rich Grassland under Long-term Climate Change.” Ecology and Evolution 7 (3): 855–62. https://doi.org/10.1002/ece3.2700.

Sigurbjörnsdóttir, Margrét Auður, Ólafur S. Andrésson, and Oddur Vilhelmsson. 2015. “Analysis of the Peltigera Membranacea Metagenome Indicates That Lichen-Associated Bacteria Are Involved in Phosphate Solubilization.” Microbiology 161 (5): 989–96. https://doi.org/10.1099/mic.0.000069.

Spribille, Toby, Veera Tuovinen, Philipp Resl, Dan Vanderpool, Heimo Wolinski, M. Catherine Aime, Kevin Schneider, et al. 2016. “Basidiomycete Yeasts in the Cortex of Ascomycete Macrolichens.” Science 353 (6298): 488–92. https://doi.org/10.1126/science.aaf8287.

Uphof, J. C. Th. 1925. “Purple Bacteria as Symbionts of a Lichen.” Science 61 (1568): 67–67. https://doi.org/10.1126/science.61.1568.67.

Vandenkoornhuyse, Philippe, Achim Quaiser, Marie Duhamel, Amandine Le Van, and Alexis Dufresne. 2015. “The Importance of the Microbiome of the Plant Holobiont.” New Phytologist 206 (4): 1196–1206. https://doi.org/10.1111/nph.13312.

Wang, Qiong, George M. Garrity, James M. Tiedje, and James R. Cole. 2007. “Naive Bayesian Classifier for Rapid Assignment of RRNA Sequences into the New Bacterial Taxonomy.” Applied and Environmental Microbiology 73 (16): 5261–67. https://doi.org/10.1128/AEM.00062-07.

Wedin, Mats, Stefanie Maier, Samantha Fernandez-Brime, Bodil Cronholm, Martin Westberg, and Martin Grube. 2016. “Microbiome Change by Symbiotic Invasion in Lichens: Microbiome Change by Symbiotic Invasion in Lichens.” Environmental Microbiology 18 (5): 1428–39. https://doi.org/10.1111/1462-2920.13032.

West, Nyree J., Delphine Parrot, Claire Fayet, Martin Grube, Sophie Tomasi, and Marcelino T. Suzuki. 2018. “Marine Cyanolichens from Different Littoral Zones Are Associated with Distinct Bacterial Communities.” PeerJ 6 (July): e5208. https://doi.org/10.7717/peerj.5208.

Xu, Maonian, Starri Heidmarsson, Margret Thorsteinsdottir, Marco Kreuzer, Julie Hawkins, Sesselja Omarsdottir, and Elin Soffia Olafsdottir. 2018. “Authentication of Iceland Moss (Cetraria Islandica) by UPLC-QToF-MS Chemical Profiling and DNA Barcoding.” Food Chemistry 245 (April): 989–96. https://doi.org/10.1016/j.foodchem.2017.11.073

